# Modular dynamics of conscious and unconscious states in marmoset cortex

**DOI:** 10.1101/2025.07.09.663972

**Authors:** Dominic I. Standage, Yuki Hori, Joseph Y. Nashed, Daniel J. Gale, Ravi S. Menon, Stefan Everling, Jason P. Gallivan

## Abstract

General anesthetics are routinely used to induce unconsciousness. While much is known about their effects on receptor function and the activity of individual neurons, much less is known about how these local effects are manifest at the level of largescale, distributed brain networks. Using functional magnetic resonance imaging (fMRI) with the common marmoset (Callithrix jacchus) we investigated the effects of the anaesthetic isoflurane on functional brain networks and their temporal dynamics, comparing network measures during wakeful rest and induced unconsciousness. The anaesthetic condition was characterised by weak functional networks that were more similar to anatomical structure and more fragmented than during wakeful rest. Conversely, the awake condition was characterised by coordinated network reconfiguration and more distinct subnetwork composition. Our findings are consistent with the view that consciousness is an emergent property of the dynamics of functional brain networks, and that anaesthetics impoverish these dynamics by reducing the efficacy of synaptic transmission.

## Introduction

Consciousness can be defined as a state of wakeful awareness, characterised by the ability to sense and respond to stimuli (see the Discussion for a less cowardly definition). Our understanding of the mechanisms underlying consciousness has been greatly informed by the study of *un*consciousness with general anaesthetics. Much of this work has focused on the cellular effects of anaesthesia (1–3), but a growing number of studies have used functional magnetic resonance imaging (fMRI) to investigate large-scale, network-level changes resulting from these cellular effects (4–6) (for review, see 7, 8). This approach assumes that consciousness is an emergent property of large-scale functional networks, the composition of which is estimated by the pairwise covariation of haemodynamic signals (9–11). The dynamics of functional networks have been shown to differ during conscious and unconscious states, with a smaller repertoire of network states observed during unconsciousness and fewer transitions between them (5). These network analyses have been invaluable for characterising the systems-level bases of consciousness, providing evidence for the impoverishment of network dynamics under anaesthetic. This approach, however, does not consider the configurations of subnetworks comprising these states, which have been shown to differ across depths of unconsciousness (6).

The brain’s functional architecture is inherently modular, comprised of densely intra-connected subnetworks with sparse connectivity between them. This organisational principle is believed to optimise distributed processing and support complex emergent dynamics (12), both of which are foundational principles of mechanistic theories of consciousness (13–15). Crucially, this organisation is not static, as functional subnetworks have been shown to assemble and disassemble over time (6, 16–18). The tracking of this reconfiguration, or *dynamic modularity*, therefore provides a window into systems-level brain function. Furthermore, metrics derived from dynamic modularity are associated with cognitive engagement (18, 19), a state intrinsically linked to conscious awareness (15, 20). We reasoned that if consciousness is an emergent property of functional brain networks, then patterns of modular reconfiguration should serve as signatures for differentiating conscious and unconscious states, potentially mirroring those observed during active cognition (18, 21).

The common marmoset (*Callithrix jacchus*) is an increasingly popular, pre-clinical primate model for investigating systems-level brain organisation (22–28). Its brain architecture is more translationally relevant to humans than that of non-primates (29, 30), an important consideration for studies of large-scale, complex brain functions such as consciousness. Protocols for resting-state fMRI under anaesthetic are well established for this species (26, 31–34), including those for isoflurane, allowing for rapid induction (of) and recovery from unconsciousness. Furthermore, extensive anatomical connectivity data from retrograde tracing and diffusion-weighted MRI are publicly available through resources such as the Marmoset Brain Connectivity Atlas (https://www.marmosetbrain.org) and the Mormoset Brain Mapping initiative (https://marmosetbrainmapping.org). The marmoset is therefore an ideal model species for comparing functional network dynamics between conscious and unconscious states.

We compared modular dynamics during consciousness (wakeful rest) and unconsciousness (isoflurane anaesthesia) with resting-state fMRI recorded from marmosets (26). We began by quantifying the strength of functional connectivity (FC) and its similarity to anatomical structural connectivity (SC) across conditions. These ‘first order’ measures have been altered by anaesthesia in previous studies (5, 35), providing an interpretive context for our primary investigation of ‘higher order’ measures of dynamic modularity (derived from FC). We then tested three principal hypotheses on the difference between conscious and unconscious states. First, we hypothesised that functional networks would be more fragmented during unconsciousness, based on evidence from studies of anaesthetics (7). We tested this hypothesis by calculating the number of dynamic modules in subjects’ functional networks in each condition, where a larger number of modules indicates greater fragmentation (6). Second, we hypothesised that the reconfiguration of functional networks would be less coordinated during unconsciousness, based on evidence linking coordinated and uncoordinated modular reconfiguration to cognition (18) and sedation (6) respectively. We tested this hypothesis by measuring the degree to which brain regions changed modular affiliation in groups and individually in each condition (36). Third, we hypothesised that the subnetwork composition of functional networks would be less clearly defined during unconsciousness, based on findings that the distinctiveness of FC subnetworks varies with the strength of sedation (6). We tested this hypothesis by calculating the stability and integrity of static ‘consensus’ subnetworks derived from the dynamic modules. These analyses served to characterise conscious and unconscious states with signatures of large-scale brain dynamics previously associated with cognitive function.

## Results

To characterise cortical modular composition and dynamics during conscious and unconscious states, we collected resting state fMRI (rs-fMRI) data from four marmosets during wakeful rest and following administration of the general anaesthetic isoflurane (26). Each marmoset’s haemodynamic signals were grouped into discrete brain regions according to the Paxinos atlas (37) and uniformly (90%) sparse functional networks were constructed from the covariation of regional activation in sliding time windows. We partitioned these ‘multislice’ networks into spatiotemporally dynamic modules and tested whether subjects’ FC was significantly modular in each condition by comparing network modularity scores to those of three classes of null model for temporal modularity: connectional, nodal and temporal null networks (38). The difference between real and null scores was significant for all three classes of model during the awake [connectional: t(3) = 15.478, p = 5.000e-4; nodal: t(3) = 21.107, p = 3.000e-4; temporal: t(3) = 6.190, p = 2.900e-3] and anaesthetic [connectional: t(3) = 4.473, p = 8.300e-3; Δ nodal: t(3) = 4.816, p = 5.200e-3; temporal t(3) = 16.891, p = 3.000e-4] conditions, where statistical significance was determined by permutation testing (see Methods). Thus, we established that participants’ intrinsic FC was composed of interacting subnetworks during consciousness and unconsciousness.

### Anaesthesia weakened functional connectivity, rendering it more similar to structural connectivity

Before testing our hypotheses on dynamic modularity, we sought to determine whether subjects’ FC would show condition-dependent changes consistent with those of earlier studies (5, 26, 35), providing a foundation for interpreting our subsequent analyses of FC dynamics. To this end, we first compared subjects’ static FC (averaged over all scans, Figure 2A) to average marmoset structural connectivity (SC) (39), measuring the Euclidean distance between FC and SC in the awake and anaesthetised conditions. As expected, the distance between FC and SC during consciousness was significantly greater than during unconsciousness (Figure 2B, see figure caption and Methods for statistics). Because (tract tracing) SC reveals monosynaptic (direct) connectivity (39–42), this finding indicates that anaesthesia reduces polysynaptic (network-level) connectivity. A such, it provides a mechanistic perspective on limitations to the brain’s functeional repertoire under anaesthetic.

Next, we measured the overall strength of FC by calculating the node strength of each region (the sum of its connection weights) and averaging across regions. Consistent with the widespread dampening effects of anaesthesia (43, 44) node strength was significantly higher during the awake condition (Figure 2C).

In summary, FC was weaker and more similar to the monosynaptic SC scaffold during unconsciousness, consistent with findings from earlier studies (5, 26, 35). These results confirm that anaesthesia produced the expected ‘first order’ effects on FC, characterised by impoverished regional interactions, setting the proverbial stage for our analysis of modular dynamics.

### Anaesthesia increased the fragmentation of functional networks

Having determined that anaesthetic-dependent changes in FC were as expected, we sought to test our hypothesis that cortical functional networks would be more fragmented during unconsciousness. Fragmentation was quantified by calculating the mean number of dynamic modules in each scan and condition (see Methods), where a higher module count corresponds to the decomposition of subjects’ FC into a larger number of smaller, segregated subnetworks. Our hypothesis was confirmed, as the number of modules was significantly greater under anaesthetic (Figure 2D). Importantly, this finding cannot be attributed to condition-dependent differences in network sparsity (inversely, density) because the spatiotemporal modules were derived from uniformly sparse networks (90% sparse in all time windows, scans and conditions). Thus, the greater number of modules under anaesthetic reveals a legitimate increase in fragmentation, directly supporting the hypothesis that anaesthetic-induced unconscsiousness results from the fragmentation of large-scale functional brain networks (6) (for review, see 7, 45) and contradicting the recent claim that such fragmentation is an artifact of statistical thresholding in network construction (46).

### Anaesthesia disrupted coordinated modular reconfiguration

To test our hypothesis that modular reconfiguration would be more coordinated during consciousness than unconsciousness, we calculated the cohesive and disjointed flexibility (a.k.a. cohesion strength and disjointedness) of subjects’ brain regions during the awake and anaesthetic conditions (see Methods). Cohesion strength refers to the number of times a given region changes modules together with each other region, summed over all other regions. In contrast, disjointedness refers to the number of times a given region changes modules on its own, relative to the number transitions (36). For each subject, scan and condition, we therefore calculated each of these measures for each brain region in each partition, before averaging all partitions, regional scores and scans (in that order).

Consistent with our hypothesis, cohesion strength was higher in the awake condition (Figure 3A) and disjointedness was higher in the anaesthetic condition (Figure 3B). Notably, these opposing signatures of network reconfiguration were evident throughout cortex, as the vast majority of brain regions showed a positive condition-dependent difference (awake minus anaesthetic) in cohesion strength (100 out of 116 regions) and a negative difference in disjointedness (100 out of 116; not the same regions) (Figure 3C). This widespread effect provides strong evidence that coordinated modular reconfiguration is a global property of cortical FC, characteristic of the conscious state and severely disrupted during anaesthetic-induced unconsciousness.

### Anaesthesia reduced the stability and distinctiveness of functional subnetworks

To test our hypothesis that FC subnetworks would be less clearly defined during unconsciousness, we identified static subnetworks that summarise the composition of the dynamic modules in the awake condition. To do so, we calculated the proportion of all modular partitions (over all subjects, scans and time windows) in which each pair of regions was placed in the same module (Figure 4A) and we clustered this module allegiance matrix (17) with symmetric non-negative matrix factorisation (see Methods). The best clustering solution identified four subnetworks (4B-D), a visuo-fronto-parietal network (blue), a ventrotemporal-prefrontal network (green), a somatomotor network (red) and an auditory network (orange).

Next, we quantified the degree of regional interaction within and between the subnetworks over time. With each brain region (for each subject) assigned to one of the four subnetworks, the interaction between any two subnetworks can be defined by *I*_*k*1,*k*2_ = (∑*i* ∈ *C*_*k*1_, *j* ∈ *C*_*k*2_*P*_*i,j*_)*/*(|*C*_*k*1_| |C_*k*2_|), where *C*_*k* ∈{1,2}_ are modules *|C_k_|* is the number of regions they contain, and *P*_*i,j*_ is the module allegiance matrix described above, quantifying the proportion of time windows in which regions *i* and *j* were in the same module. We refer to the interaction of a subnetwork with itself as recruitment, calculated by allowing *k*1 = *k*2. The integration between two subnetworks *k*1≠ *k*2 is the normalised interaction between them 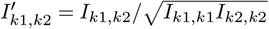 (17). In effect, recruitment captures the stability of a subnetwork’s membership over time, whereas integration captures the interactions between subnetworks.

Both metrics revealed significant differences between conditions. Recruitment was weaker in three out of four subnetworks in the anaesthetic condition (all except the auditory subnetwork; Figure 5A). Conversely, integration *between* subnetworks was significantly greater under anaesthetic for all four subnetworks (Figure 5B). Together, these findings confirm our hypothesis, showing that functional networks become less stable and more ‘smeared’ during unconsciousness, losing their compositional distinction relative to consciousness.

## Discussion

We investigated the effects of the general anaesthetic isoflurane on cortical functional networks and their modular dynamics in the common marmoset, comparing network measures during wakeful rest and induced unconsciousness. We began by establishing that anaesthesia weakened functional networks (Figure 2C) and rendered FC more similar to SC than during wakefulness (Figure 2B). These findings are consistent with those of earlier studies (5, 26, 35) and they identify the background conditions for impoverished network dynamics during unconsciousness. Our primary analysis then tested and confirmed three hypotheses on these dynamics. First, we detected more dynamic modules in the anaesthetic condition (Figure 2D), confirming the hypothesis that FC would be more fragmented during unconsciousness. Second, we found that modular flexibility was less cohesive (Figure 3A) and more disjointed (Figure 3B) under anaesthetic, confirming the hypothesis that modular reconfiguration would be less coordinated during unconsciousness. Third, we found that functional subnetworks derived from the dynamic modules (Figure 4) showed weaker recruitment (Figure 5A) and greater integration (Figure 5B) under anaesthetic, confirming the hypothesis that subnetwork composition would be less clearly defined during unconsciousness. Together, our findings demonstrate that anaesthetics impoverish the rich dynamic repertoire of awake brain, supporting the view that consciousness is an emergent property of these dynamics.

### A cascade of mechanisms from impaired synaptic function to disrupted network dynamics

Our results point to a cascade of mechanisms initiated by the dampening effect of anaesthetics on synaptic transmission (43, 44). This impairment of synaptic function is presumably the cause of the reduction in overall FC strength (Figure 2C) that renders FC more similar to its underlying anatomical structure (Figure 2B). Combined with the diminishment of sustained longrange coupling (26), this regression to the structural connectome effectively divides functional networks into a larger number of smaller modules (Figure 2D). Importantly, it also reflects limitations on the brain’s dynamic repertoire, quantified here by less coordinated, more disjointed network reconfiguration (Figure 3) and a pronounced reduction in the stability and relative segregation of subnetwork composition (Figure 5).

Whether FC fragmentation is the cause of unconsciousness or merely one of its properties is a matter of interest. Notably, our use of uniformly sparse networks demonstrates that fragmentation is not an artifact of network sparsity, as claimed by an earlier study (46). The underlying premise of this claim is that weaker FC entails greater sparsity when a statistical threshold is used in the estimation of functional networks, and that sparser connectivity is inherently more fragmented. For example, if FC connections (correlations between regional timeseries) with a corresponding p-value less than 0.05 are set to 0, then weaker FC must also be sparser. While our study shows that this premise does not explain away fragmentation as a neural correlate of unconsciousness, we cannot claim that fragmentation causes unconsciousness. Nonetheless, our findings are consistent with a large body of data providing evidence for mechanistic theories of consciousness that relate to fragmentation (7). In particular, integrated information theory (IIT) (13) and global neuronal workspace theory (GNWT) (14, 47) emphasise the capacity for information integration and global availability of information, respectively, predicting that the disruption and discoordination of large-scale networks will impair consciousness. The fragmentation (Figure 2D), discoordination (Figure 3) and loss of subnetwork distinctiveness (Figure 5) shown here confirm these predictions, potentially reflecting impaired information integration (per IIT) and/or the failure to establish a stable global workspace (per GNWT).

#### Theoretical implications for hierarchical processing

These theoretical accounts of consciousness point to the importance of the brain’s hierarchical structure. Association cortices sit atop this hierarchy (48, 49) and are central to integration under IIT and to a global workspace under GNWT (7). Accordingly, anaesthesia should be expected to disproportionately disrupt connectivity with and among higher-order brain regions (7, 14), but the subnetwork disintegration found here (Figure 5A) did not show a hierarchical effect. This finding contrasts with previous, anaesthetic dose-dependent observations in macaques, where the integrity of a sensorimotor network was spared in comparison to higher-order networks (6). A possible explanation for this discrepancy is that the single, standard (26) anaesthetic dose used here (1.5% isoflurane) induced sufficiently widespread synaptic impairment (43, 44) and thalamocortical disruption (26) to dominate hierarchical effects. Alternatively, the lack of hierarchical selectivity may reflect the neurobiology of the marmoset itself (27). The functional segregation of association networks and sensorimotor networks, as well as the dorsal and ventral processing streams, may be less pronounced in the common marmoset than in macaques and humans, potentially relating to differences in patterns of cortical expansion during primate evolution (27, 50). If the operational distinctions between such large-scale systems are less pronounced in marmosets, a global perturbation like general anaesthesia may produce more uniform effects across these systems. Future studies varying anaesthetic dose (*e.g.* 6) with marmosets are critical for assessing these possibilities. Additionally, dynamic network analyses across primate species would quantify the extent to which hierarchical distinctions vary and how this variation influences susceptibility to anaesthetic agents (27, 29).

### From sensation and responsiveness to subjective experience

Our use of general anaesthesia with the common marmoset necessarily limits our operational definition of consciousness and the scope of our study to the ability to sense and respond to stimuli, as noted in the Introduction. Nonetheless, our hypotheses and findings have implications for the more general (and philosophically satisfying) phenomenon of experiential consciousness. Sensation and responsiveness have qualitative characteristics. In Nagel’s phrasing (51), there is ‘something that it is like’ to sense and respond to stimuli. These subjective qualities [a.k.a. qualia (52)] are central to what has been called the hard problem of consciousness (53) and warrant some discussion (see also 15). Of course, this discussion requires us to assume a degree of generalisation from marmosets to humans, given that marmosets’ cannot linguistically report their experiences. In the context of FC dynamics, this assumption appears to be justified, as our predictions for marmosets were directly informed by the whole-brain characterisation of human cognition with the same analysic methods (18, 21) and were subsequently confirmed by our results. Additionally, studies have demonstrated strong functional correspondence between specific brain regions in humans and marmosets (29), supporting the translational relevance of our findings.

The phenomenon of dreaming may provide the most instructive distinction between ‘the ability to sense and respond to stimuli’ and the notion of subjective experience. Sleep is conventionally considered an unconscious phenomenon, but we subjectively experience our dreams (54, 55) (see 56 for a more nuanced description). This juxtaposition not only highlights a point of philosophical interest, but it also makes three specific predictions based on our findings: during rapid eye movement (REM) sleep, FC will be more fragmented, and will show more coordinated reconfiguration and less clearly defined subnetwork composition than during nonREM sleep. These predictions could be tested by running our modularity analysis with sleeping human participants, measuring the number of detected modules (Figure 2D), their cohesive and disjointed flexibility (Figure 3), and the recruitment and integration of subnetworks derived from module allegiance matrices (Figure 5). fMRI is quite commonly used with non-anaesthetised, sleeping human participants (56–58) and future studies should test our predictions by analysing such data.

The possibility of machine consciousness is a second point of interest. The current excitement (and hyperbole) in the field of artificial intelligence (AI) has lead to significant public interest in this topic, as AI enthusiasts ask whether conscious experience is exclusive to animal species (most notably humans) or whether it could be supported by inorganic machinery. In our view, large artificial neural networks (ANN) with dense feedback connectivity could, in principle, support consciousness. However, the ANNs that have captured the public imagination (*e.g.* large language models built on ‘transformer’ architectures) are almost exclusively feed-forward, *i.e.* their connectivity propagates activity in one direction only, from input to output. This architectural consideration points to two related discussion points. Firstly, feed-forward networks with at least one hidden layer (often called deep networks) are universal function approximators, that is, they can approximate any input-output mapping with arbitrary accuracy (given enough training time and a sufficiently large hidden layer) (59). Thus, if consciousness requires the emergent properties of feedback dynamics (14) (potentially revealed by the coordinated reconfiguration of functional networks, per our findings here) then consciousness is more than an inputoutput mapping. Secondly, most (perhaps all) mechanistic theories of consciousness include some kind of feedback processing, implying that feed-forward networks cannot be conscious. These theories include IIT (13), GNWT (14, 47), dendritic integration theory (60) and, unsurprisingly, ‘recurrent’ hypotheses on consciousness (61, 62). IIT, in particular, has several fascinating implications for machine consciousness, including the implication that a faithful digital *simulation* of consciousness cannot actually be conscious, and that a feedforward network that is formally equivalent to a conscious recurrent network cannot itself be conscious (63). Sorry, chatGPT. For a thorough treatment of theoretical frameworks for consciousness, see Seth and Bayne 15.

The use of ANNs to characterise neural circuit motifs and their computations is an active methodology of systems neuroscience (64–66), providing insights into the mechanistic bases of perception (67, 68), cognition (69, 70) and action (71, 72). It is not obvious how these methods can best be used to study consciousness, since the guiding principle of this ‘neuro-AI’ approach is to use machine learning algorithms to train a network to perform a challenging task and then to report features and processing principles of the trained network. To us, it seems unlikely that consciousness is something that can be trained by the optimisation of an objective function (though see 73 for some ideas). In our view, a more promising approach is to simulate brain circuitry (characterised by dense recurrent connectivity) with neural models and to compare model activity with neural recordings from conscious and unconscious states (14, 74). This approach serves to characterise consciousness by its neural signatures across resolutions, scales and species, making testable predictions for experiments. From a scientific research perspective (as opposed to a philosophical one), we cannot think of a more illustrative example of what makes ‘the hard problem’ so hard: there may be no clear behavioural signature of subjective experience. Fortunately, there is a growing body of evidence for neural signatures of (un)consciousness (7, 14, 45, 60, 74–77) including those described by our study. In this regard, a promising direction in neuroimaging and neural modelling is the fitting of SC models to FC data, characterising the dynamics of state transitions and identifying circuit mechanisms that support them (78, 79). Future work should apply this methodology to the study of consciousness.

### Methodological considerations and future directions

Within the framework of dynamic modularity, we have revealed significant differences in the spatiotemporal organisation of cortical FC between conscious and anaesthetised states in marmosets. Nonetheless, several considerations warrant discussion and inform future research directions. Firstly, our findings are specific to 1.5% isoflurane following ketamine induction (26). While this approach is standard, it calls for caution in generalising our results. Different anaesthetic agents have distinct molecular targets and can influence network connectivity in different ways (80, 81). For example, isoflurane significantly attenuates thalamic and interhemispheric FC in marmosets (26). Determining the generality of the network effects reported here requires the systematic variation of anaesthetic dose (e.g. 6) across different agents. Such work is crucial for isolating the fundamental mechanisms of unconsciousness from the effects of specific agents, potentially augmented by alternative protocols that may preserve function connectivity, such as the combination of low-dose medetomidine and isoflurane (26, 82). The residual effects of ketamine induction are an additional factor in this context (46).

Secondly, understanding the effects of anaesthetics across the cortical hierarchy remains a key challenge. Our finding of relatively uniform subnetwork disintegration contrasts with earlier work with macaques showing the relative sparing of sensorimotor connectivity across levels of anaesthetic dose (6). As touched upon above, the uniform effect shown here may reflect anaesthetic dose or inter-species differences in the organisation of large-scale functional networks. In combination with comparative studies across species (27, 29), future analyses of directed (83) and laminar (84) connectivity are required for a more comprehensive understanding of the mechanisms by which anaesthesia disrupts information flow across the cortical hierarchy.

Thirdly, complementary methods would provide further insights into the whole brain dynamics of consciousness. Explicitly estimating discrete whole-brain FC states and their transition dynamics would be a valuable next step (*e.g.* (85, 86). This approach would complement our modularity analysis by characterising the repertoire, persistence and sequencing of brain states and their transitions, offering a different lens on the impoverished dynamics reported here. We anticipate that such measures will provide additional evidence for theories linking consciousness to the to flexible traversal of a rich space of brain states (14, 87).

Finally, like all fMRI studies, ours is based on the blood oxygenation-level dependent (BOLD) signal, an indirect measure of neural activity with limited temporal resolution. While dynamic FC methods extract valuable information from these signals, integrating this approach with electrophysiological recordings (*e.g* electroencephalogram and local field potentials) may clarify the ways in which haemodynamic networks reflect real-time neural processing and network dynamics in relation to consciousness.

### Summary and conclusions

We subscribe to the view that consciousness is an emergent property of distributed brain dynamics (14, 45, 74). Using rs-fMRI with the common marmoset, our study provides a novel characterisation of brain dynamics during conscious and anaesthetic-induced unconscious states. This characterisation leverages methods that have been informative about human cognition (17–19, 21), providing an interpretive framework for our hypotheses and results that in turn leverages the conceptual, computational and functional overlap between conscious awareness and cognitive engagement (15, 20). Our results point to coordinated reconfiguration of relatively stable, distinct subnetworks as a signature of the conscious brain. This signature is in stark contrast to the weak, fragmented connectivity, poorly defined subnetwork composition, and uncoordinated dynamics of the anaesthetised brain. As such, our study demonstrates that brain dynamics are severely impoverished during anaesthesia-induced unconsciousness, providing evidence that consciousness emerges from the rich dynamic repertoire of the awake brain.

## Methods

### Overview

To characterize the effects of general anesthesia on the modular dynamics of cortical functional networks, we analysed a previously published resting-state functional magnetic resonance imaging (rs-fMRI) dataset acquired from common marmosets (*Callithrix jacchus*) at 9.4 Tesla MRI (26). The dataset enabled the comparison of large-scale brain network organization between conscious wakeful rest and unconsciousness induced by the administration of 1.5% isoflurane. Following preprocessing of the functional data (detailed below), each animal’s cerebrum was parcellated into discrete cortical regions based on the Paxinos atlas (37). Regional BOLD time series were extracted and dynamic FC matrices were constructed by computing pairwise correlations within sliding temporal windows. These ‘multislice’ networks were partitioned into dynamic modules by a time-resolved clustering (a.k.a. community detection) algorithm to quantify their reconfiguration over time (38). We then compared measures of network fragmentation and modular dynamics in the awake and anaesthetised conditions.

### Animal preparation

All experimental procedures involving animals were conducted in strict accordance with the guidelines set forth by the Canadian Council of Animal Care and received prior approval from the Animal Care Committee at the University of Western Ontario. The study included four common marmosets (*Callithrix jacchus*), one female and three male, with an average weight of 323 *±* 61 g and an average age of 1.6 *±* 0.3 years at the commencement of the experiments (26). Prior to MRI scanning, each marmoset underwent an aseptic surgical procedure to implant a head chamber, which served to immobilize the head during image acquisition, following established protocols (88). Briefly, animals were placed in a stereotactic frame (Narishige Model SR-6C-HT) and the 3D-printed head chamber (Formlabs Clear Resin V4, 0.25 mm resolution) was affixed to the skull using adhesive resin (All-bond Universal Bisco) and dental cement (C & B Cement, Bisco), ensuring precise positioning and orientation.

### Imaging hardware and acquisition parameters

Magnetic resonance imaging was performed using a 9.4 Tesla, 31-cm horizontal bore scanner (Varian/Agilent) controlled by a Bruker BioSpec Avance III console operating with Paravision-6 software (Bruker BioSpin Corp). This system included a specialized, high-performance 15-cm diameter gradient coil capable of achieving a maximum strength of 400 mT/m (89). Radiofrequency transmission utilized a 12-cm inner diameter quadrature birdcage coil, while signal reception was accomplished using a custom-designed 5-channel receiving coil optimized for the marmoset head (90). The imaging experiments were conducted at the Centre for Functional and Metabolic Mapping within the Robarts Research Institute at The University of Western Ontario.

Functional images were acquired using a single-shot gradient-echo echo-planar imaging (GE-EPI) sequence, using the following parameters: repetition rime (TR) = 1500 ms, echo time (TE) = 15 ms, flip angle = 40^°^, field of view (FOV) = 64 *×* 64 mm, matrix size = 128 *×* 128, resulting in an isotropic voxel size of 0.5 mm. Forty-two contiguous slices covered the brain volume. The acquisition bandwidth was 500 kHz, and parallel imaging (GRAPPA) with an acceleration factor of 2 in the anterior-posterior direction was employed. For each animal and condition, six functional runs were acquired, each consisting of 600 volumes, corresponding to 15 minutes of scan time per run. Additionally, a high-resolution T2-weighted anatomical reference image was obtained for each animal using a rapid acquisition with refocused echoes (RARE) sequence (TR = 5500 ms; TE = 53 ms; FOV = 51.2 *×* 51.2 mm; matrix size = 384 *×* 384; voxel size = 0.133 *×* 0.133 *×* 0.5 mm; 42 slices; bandwidth = 50 kHz; GRAPPA acceleration factor = 2).

### Animal training

To prepare marmosets for awake imaging, a multi-week acclimatisation and training regimen was implemented, adapted from established methods (91). This process began with habituating the animals to restraint within a specialised holder using neck and tail plates for progressively longer durations (up to 30 minutes). Subsequently, the restraint tube was placed inside a mock MRI scanner bore, simulating the scanning environment while exposing the animals to recorded scanner sounds at increasing volumes (up to 80 dB) for extended periods (up to 60 minutes). In the final training phase, animals were head-fixed using their implanted chambers and fixation pins within the mock scanner, again with exposure to MRI sounds. Throughout training, animals received positive reinforcement (*e.g.* pudding, marshmallow fluff) for maintaining a calm, forward-facing posture with minimal limb movement. Animal tolerance and calmness were systematically evaluated using a behavioral rating scale (91). All marmosets included in the study successfully completed the training, achieving scores indicative of quiet, calm behavior suitable for awake imaging.

### Image acquisition

#### Image acquisition in the awake condition

For the awake scanning sessions, the trained marmosets were carefully placed and secured in the animal holder using the neck and tail plates. Inside the MRI scanner room, their heads were firmly fixed using the fixation pins connected to the implanted chamber, a procedure designed to minimize the time the animal spent head-fixed prior to scanning (34). This head-fixation system effectively minimizes head motion during data acquisition. To mitigate magnetic susceptibility artifacts near the brow ridge, a lubricating gel (MUKO SM321N) was applied within the chamber before scanning commenced (34). Six functional runs were acquired per awake session.

#### Image acquisition in the anaesthetised condition

Anesthetised scanning sessions took place a minimum of one week after the awake sessions to avoid potential confounding effects of anesthetic agents on cerebrovascular physiology (92). Anesthesia was initially induced using an intramuscular injection of ketamine (20 mg/kg). Once positioned in the animal holder identically to the awake setup, anesthesia was maintained throughout the scan via inhalation of 1.5% isoflurane delivered through a custom-made anesthesia mask (34). The carrier gas for isoflurane was medical air (21% *O*_2_). Vital signs, including respiration rate, peripheral oxygen saturation (SpO2), and heart rate, were continuously monitored using pulse oximetry by a veterinary technician and maintained within normal physiological ranges. Core body temperature was regulated using a combination of warm-water circulating blankets, thermal insulation materials, and warmed air. Six functional runs were acquired per anesthetised session.

### Image preprocessing

Anatomical T2-weighted images underwent preprocessing primarily for accurate brain extraction. An initial mask was generated using FSL’s (93) Brain Extraction Tool (BET), followed by registration of the National Institutes of Health (NIH) marmoset T2w template (37) to the individual’s T2w scan using FMRIB’s linear (FSL FLIRT) and non-linear (FSL FNIRT) alignment algorithms. This refined registration allowed for the creation of a precise individual brain mask used for subsequent steps.

Functional MRI data preprocessing was performed on the raw image files following internal quality control assessments. The preprocessing pipeline was as follows: raw data were first converted to the Neuroimaging Informatics Technology Initiative (NIfTI) format and organized according to the Brain Imaging Data Structure (BIDS) standard (94). The initial four volumes from each functional run were discarded to ensure magnetization equilibrium. Head motion was corrected within each run using FSL MCFLIRT. The previously derived individual brain masks were applied to remove nonbrain tissue. Functional images were spatially smoothed with a Gaussian kernel of 1.0 mm FWHM. Crucially, temporal filtering was applied using a high-pass filter set to 0.01 Hz to remove low-frequency noise and scanner drift; no low-pass filter was applied. Nuisance variance was removed from the smoothed data via linear regression. The confound model included 24 motion parameters derived from the motion correction step (6 rigid-body parameters, their first temporal derivatives, and the squares of all 12 values), along with the mean time series extracted from white matter (WM) and cerebrospinal fluid (CSF) compartments. Finally, the fully preprocessed functional data for each run were registered to the standard NIH marmoset T2w template space (37) using FSL FLIRT and FNIRT.

The preprocessed fMRI images were then parcellated into 116 cortical regions using the digital Paxinos marmoset atlas (37, 95), which integrates the corresponding left and right hemispheric regions (34). For each region, we computed a single time series by averaging the preprocessed BOLD signal across all constituent voxels.

### Static functional connectivity and structural comparison

To establish baseline network characteristics and compare functional organisation to underlying anatomy, we computed static functional connectivity (FC) matrices for each subject and condition. These were generated by calculating the Pearson correlation coefficient between the full, preprocessed BOLD time series for every pair of the 116 cortical regions. For comparison, we utilized a representative structural connectivity (SC) matrix for the common marmoset. This SC matrix was derived from the integrated resource described by Tian et al. 39, which combines data from sources including diffusion-weighted imaging and tract-tracing experiments. We normalised the SC matrix for comparison with FC by dividing all connection weights by the maximum value in the matrix, resulting in weights ranging from 0 to 1. To quantify the similarity between the observed static FC (averaged across subjects within each condition) and the above SC matrix we calculated the Euclidean distance between the flattened upper triangle of the matrices.

#### Temporal modularity of functional networks

For dynamic FC analysis, each of the 116 regional time series was divided into windows of 90 seconds (60 imaging volumes) where contiguous windows overlapped by 45 seconds, *i.e.* by 50% (6, 96). We constructed functional networks in each time window by calculating the Pearson correlation coefficient between each pair of brain regions (Figure 1C) and retained the strongest 10% of these functional connections (all other connections were set to zero). Thus, FC was 90% sparse in all time windows for all subjects. This approach served to mitigate against the possibility of spurious connectivity, while ensuring uniformly sparse FC, a requirement for comparing node strength in each condition (Figure 2C) and for addressing the claim that fragmentation of FC during unconsciousness is an artifact of non-uniform network sparseness (46). We determined the modular composition of the resulting multislice networks with a generalised Louvain method for time-resolved clustering (97). This algorithm was repeated 100 times, resulting in 100 clustering solutions (a.k.a. partitions), each of which maximised a modularity ‘quality’ function (38). On each repetition, we initialised the algorithm at random (uniform distribution) and iterated until a stable partition was found, *i.e.* the output on iteration *n* served as the input on iteration *n* + 1 until the output matched the input (16, 38). Nodes were assigned to modules at random from all assignments that increased the quality function, and the probability of assignment was proportional to this increase. We used the standard spatial and temporal resolution parameters *γ* = 1 and *ω* = 1 respectively, and the null model for positive and negative edges by Rubinov and Sporns (98) for the quality function (12, 38).

**Fig. 1.**
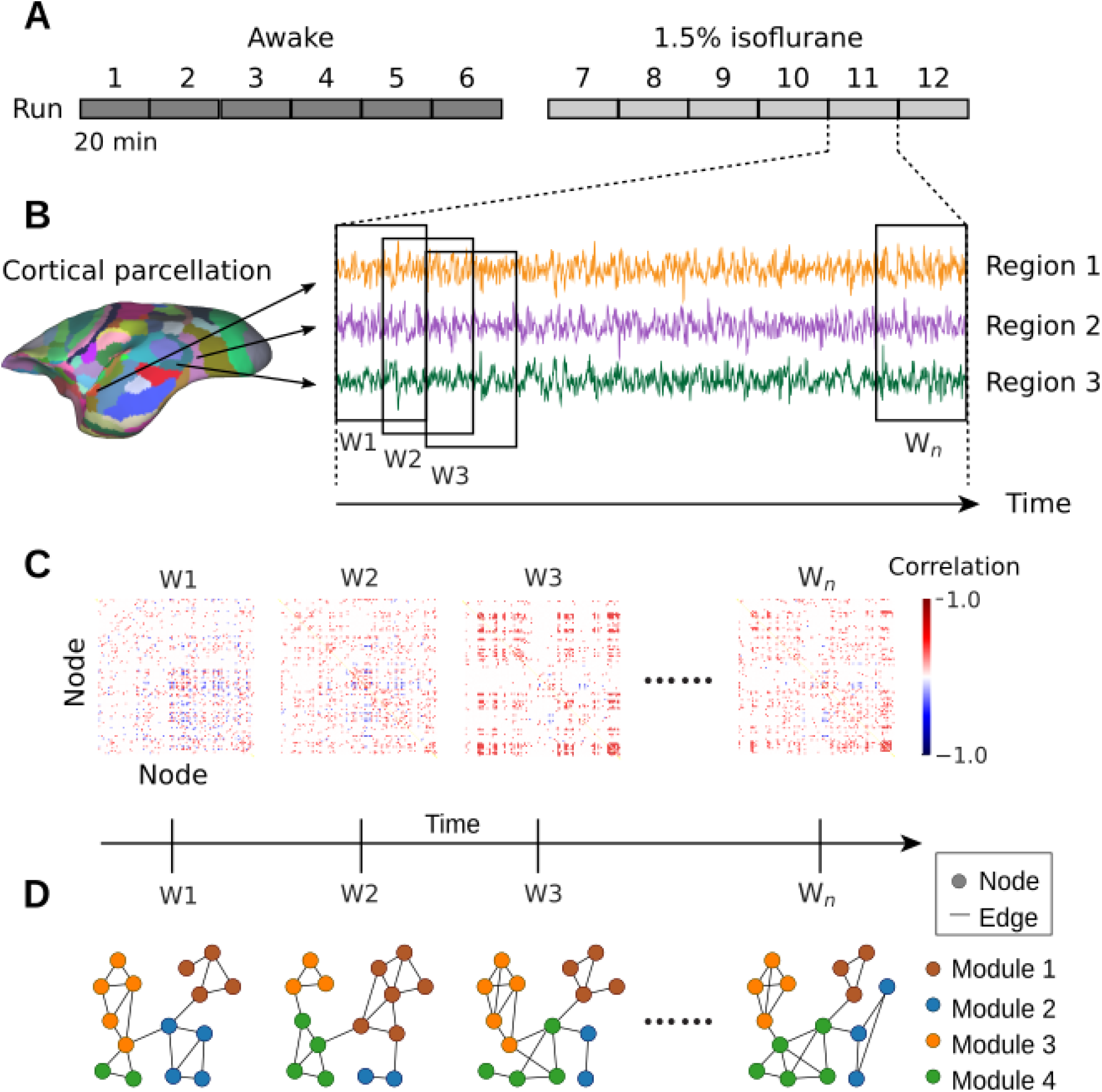
Overview of the experiment and analysis. **(A)** For each animal, six 20-minute scans were recorded during wakeful rest and under anaesthetic (1.5% isoflurane dose). **(B)** Voxel-wise fMRI signals from each animal’s cerebrum were assigned to 116 discrete regions according to the Paxinos brain atlas (37), and the average BOLD time series was extracted from each region (three example regions shown in the schematic). **(C)** The Pearson correlation coefficient was calculated for each pair of regions in sliding, half-overlapping time windows, resulting in functional connectivity (FC) matrices for each window (W1 — W_*n*_). Together, these FC matrices constitute a multi-slice (here temporal) network for each subject and scan. **(D**) Time-resolved clustering methods were applied to the multi-slice networks, identifying dynamic modules across time slices (4 modules in the schematic).

To determine the significance of temporal modularity, we compared quality function scores to those of three classes of null model known as connectional, nodal and temporal null networks (16, 38). Connectional null networks are constructed by scrambling the connectivity within each slice of a multislice network (*i.e.* within each time window), nodal null networks are constructed by scrambling the connectivity between slices, and temporal null networks are constructed by scrambling the order of slices. For each class of null network, we generated 100 partitions in the same way as for the real networks, calculating the mean quality function score (over the partitions) before taking the mean over scans for each marmoset. Statistical significance was determined by permutation testing against the null hypothesis that there is no difference in modularity between real and null networks. For each subject, the mean (over the six scans) was calculated for each of the real and null networks and then the tstatistic was calculated across subjects 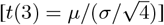. The null distribution was generated by randomly labelling six of the twelve scans as real and six as null (within each subject) and then calculating the t-statistic in the same way as for the unpermuted data. This procedure was repeated 10,000 times and the difference between real and null networks was considered significant if the unpermuted t-statistic was in the 95th percentile of the distribution.

#### Module allegiance matrix

We constructed a matrix *T*, where the elements *T*_*i,j*_ refer to the number of times regions *i* and *j* were assigned to the same module over all scans, time slices, partitions and subjects in each of the awake and anaesthetic conditions. We then constructed the module allegiance matrix *P* = (1*/C*)*T*, where *C* is the total number of time slices in all of these partitions. Values that did not exceed those of a null model were set to 0 (19).

#### Clustering of the module allegiance matrix with non-negative matrix factorisation

To derive summary networks from the module allegiance matrix, we used symmetric nonnegative matrix factorisation [SymNMF; (99, 100)], which decomposes a symmetric matrix *Y* as *Y* = *AAT*, where *A* is a rectangular matrix containing a set of non-negative factors that minimise a sum-of-squares error criterion. SymNMF has previously shown good performance in community detection problems (100, 101) and has the benefit of producing a natural clustering solution by assigning each observation to the factor on which it loads most strongly. We used the Newton-like SymNMF algorithm described by Kuang et al. (100), where for each number of factors between 2 and 10, we fit the model using 250 random initialisations of the factor matrix H, with entries sampled from a uniform distribution on the interval [0, 1]. For each rank, we calculated the average root mean squared reconstruction error, the dispersion coefficient (102) and the cophenetic correlation (103). The latter two measures quantify the consistency of the clustering solutions returned by different initialisations. We judged that a rank 4 solution was acceptable, as it explained over 85% of the variance in the observed data, it was acceptable by both the dispersion and cophenetic correlation criteria, and the generation of additional clusters provided little improvement in explained variance (Figure 4B). We then selected the rank 4 solution with the lowest reconstruction error, and generated a clustering solution by assigning each brain region to the factor with the highest loading.

### Dynamic network metrics

Following dynamic modularity detection and subnetwork identification, we quantified fragmentation and modular dynamics (respectively) in the awake and anaesthetic conditions.

#### Number of modules

To quantify network fragmentation (Figure 2D) on each scan, we calculated the number of modules detected within each partition, for each of the 100 repetitions of the generalised Louvain algorithm. The mean number of modules per scan was then calculated across partitions for each subject and condition. Given that uniformly sparse networks were used, differences in the number of modules reflect genuine changes in network fragmentation rather than thresholding effects (46).

**Fig. 2.**
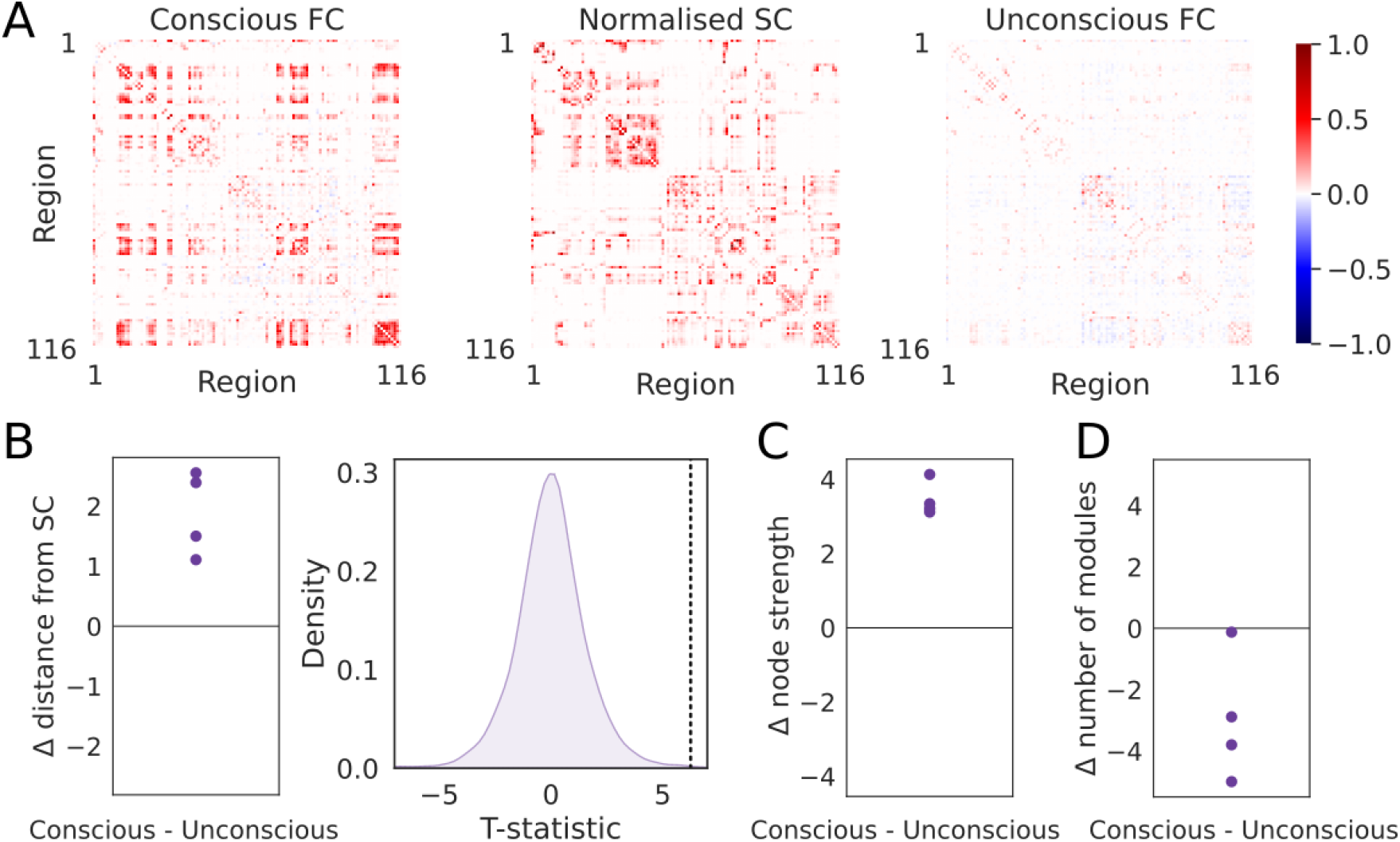
Static functional connectivity differed between conscious and unconscious states and showed established effects of anaesthesia. **(A)** FC matrices during consciousness (left) and unconsciousness (right), averaged over scans and subjects. Structural connectivity (SC) is shown between FC plots, derived from diffusion-weighted data (37) and normalized to range from 0 to 1. FC was calculated with Pearson’s correlation coefficient. All matrices use the same scale and color scheme (inset color bar). **(B)** Consistent with earlier work (5), the Euclidean distance between FC and SC was significantly greater [t(3) = 6.255, p = 0.004] during consciousness than unconsciousness, shown as the difference Δ between the awake and anaesthetic conditions. Data points correspond to subjects. Statistical significance was determined by permutation testing against the null hypothesis that there is no difference between conditions, where the t-statistic corresponding to the real difference (dotted vertical line) was in the 95th percentile of the distribution of permuted t-values (kernel density estimate, right. See Methods). **(C)** Consistent with earlier work (5, 35), overall FC strength was significantly greater during consciousness than unconsciousness [t(3) = 17.254, p = 2.000e-4], measured by node strength (sum of connection weights per region, averaged across regions). Plotting conventions are as in B. **(D)** FC was more fragmented during unconsciousness than during consciousness, as the mean number of modules was significantly greater in the anaesthetic condition than the awake condition [t(3) = −3.288, p = 0.017], quantifying condition-dependent fragmentation.

#### Flexibility metrics: cohesion strength and disjointedness

To quantify the coordination of modular reconfiguration (Figure 3), we calculated the cohesive and disjointed flexibility of each brain region (6, 36). Cohesive flexibility (a.k.a. cohesion strength) quantifies the degree to which a region changes modular affiliation *together* with other regions from its current module, across adjacent time windows. Thus, for each region *i* and all transitions between time windows *t* and *t* + 1, cohesion strength is the sum over all other regions *j* of the number of times regions *i* and *j* were in the same module in window *t* and remained together in a different module in window *t* + 1, divided by the number of transitions between windows (the number of time windows minus one). Disjointed flexibility (a.k.a. disjointedness) quantifies the degree to which a region changes modular affiliation *independently* of other regions, *i.e.* modular transitions without other regions from its current module. Thus, for each region *i*, disjointedness is the number of times region *i* changes modular affiliation between time windows *t* and *t* + 1 while no other region in its module at time *t* makes the same transition, divided by the number of transitions (36). For each subject and scan, these measures of modular flexibility were calculated for each region in each of the 100 partitions, and then averaged across partitions, regions and finally scans within each condition.

**Fig. 3.**
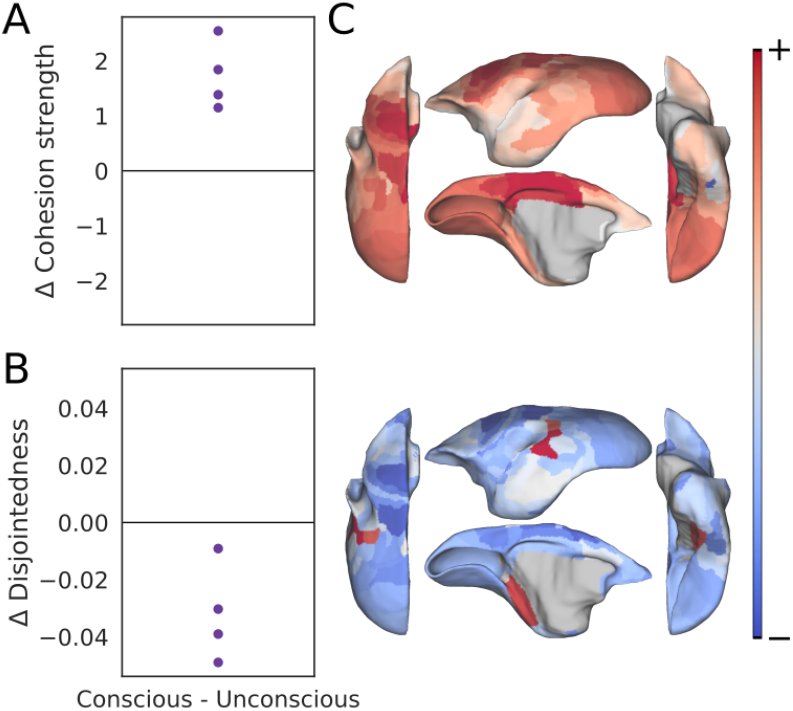
Anaesthesia disrupted coordinated of modular reconfiguration, decreasing cohesion strength and increasing disjointedness. **(A)** Mean cohesive flexibility (cohesion strength, see text) was greater during consciousness than unconsciousness [t(3) = 6.502, p = 0.005], shown as the difference Δ between the awake and anaesthetic conditions. **(B)** Mean disjointed flexibility (disjointedness, see text) was greater during unconsciousness than consciousness [t(3) = −4.338, p = 0.008] **(C)** Brain plots show difference (awake minus anaesthetic) in cohesion strength (upper) and disjointedness (lower) between conditions for each brain region. Condition-dependent differences in cohesion strength (mostly positive) and disjointedness (mostly negative) are shown with a divergent color scheme, ranging from strongly negative (dark blue) to strongly positive (dark red). For clarity, both measures were normalised before plotting.

**Fig. 4.**
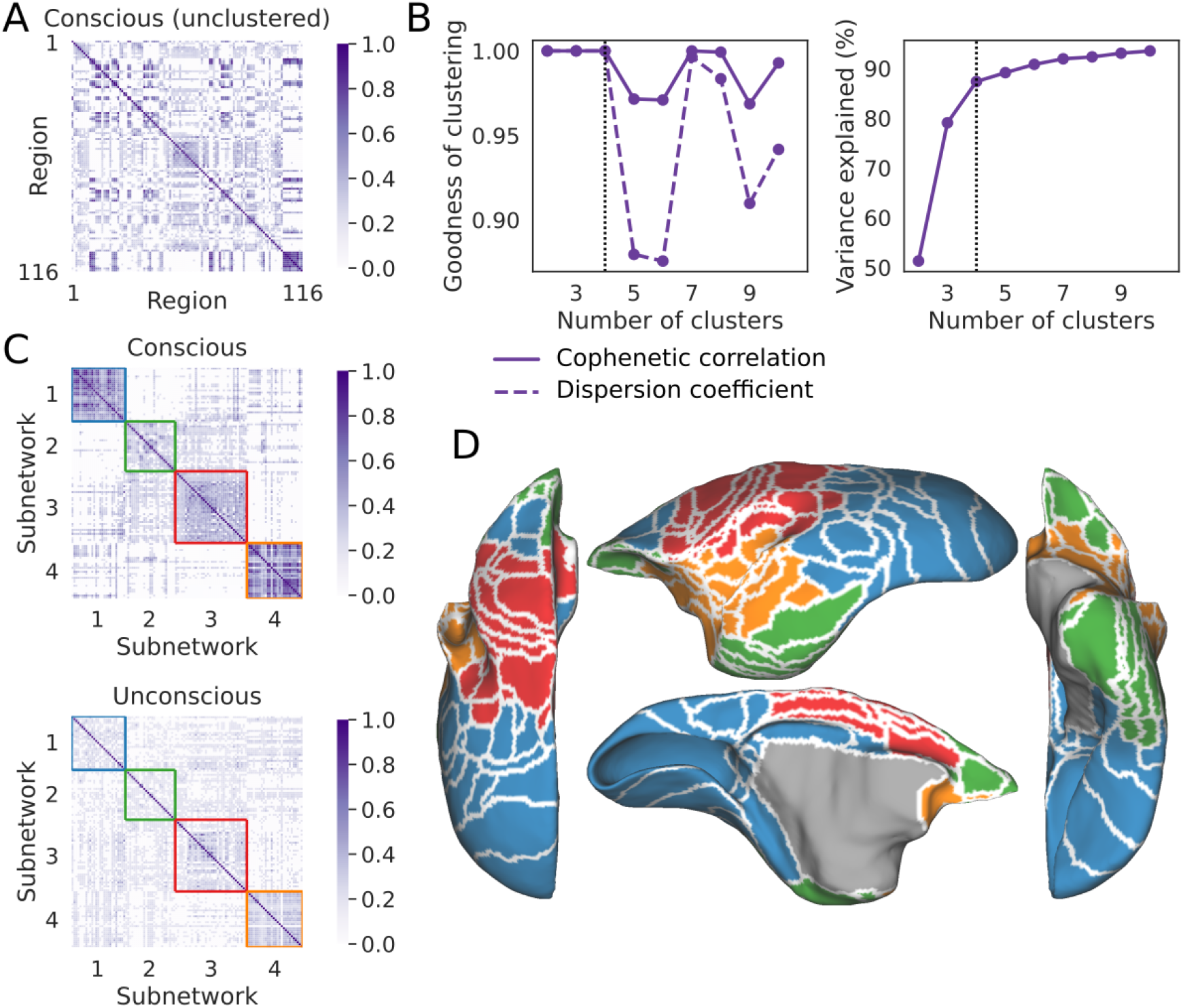
Static ‘consensus’ subnetworks derived from dynamic modules. **(A)** Module allegiance matrix from the awake condition, quantifying the proportion of modular partitions in which each pair of brain regions was in the same module. Darker shades indicate that regions frequently cluster together (see colour bar). **(B)** The module allegiance matrix shown in panel *A* was clustered with symmetric non-negative matrix factorisation for 2 to 10 factors. A 4-cluster solution was chosen (dotted vertical line in each plot), as it maximized indices of clustering stability (cophenetic correlation and dispersion coefficient, left plot), explained substantial variance in the data (85%) and showed a prominent ‘elbow’ in the explained variance, beyond which the explained variance showed incremental improvement (right plot). **C** Module allegiance matrix in the awake (conscious, top) and anaesthetised (unconscious, bottom) conditions, where brain regions are re-ordered according to the 4-subnetwork clustering solution derived from the awake data. Coloured squares outline the identified consensus subnetworks. **D)** Cortical surface maps showing the spatial topography of the four subnetworks: the visuo-fronto-parietal network (blue), the ventrotemporal-prefrontal network (green), the somatomotor network(red) and the auditory network (orange). Colours correspond to the outlined squares in panel *C*.

**Fig. 5.**
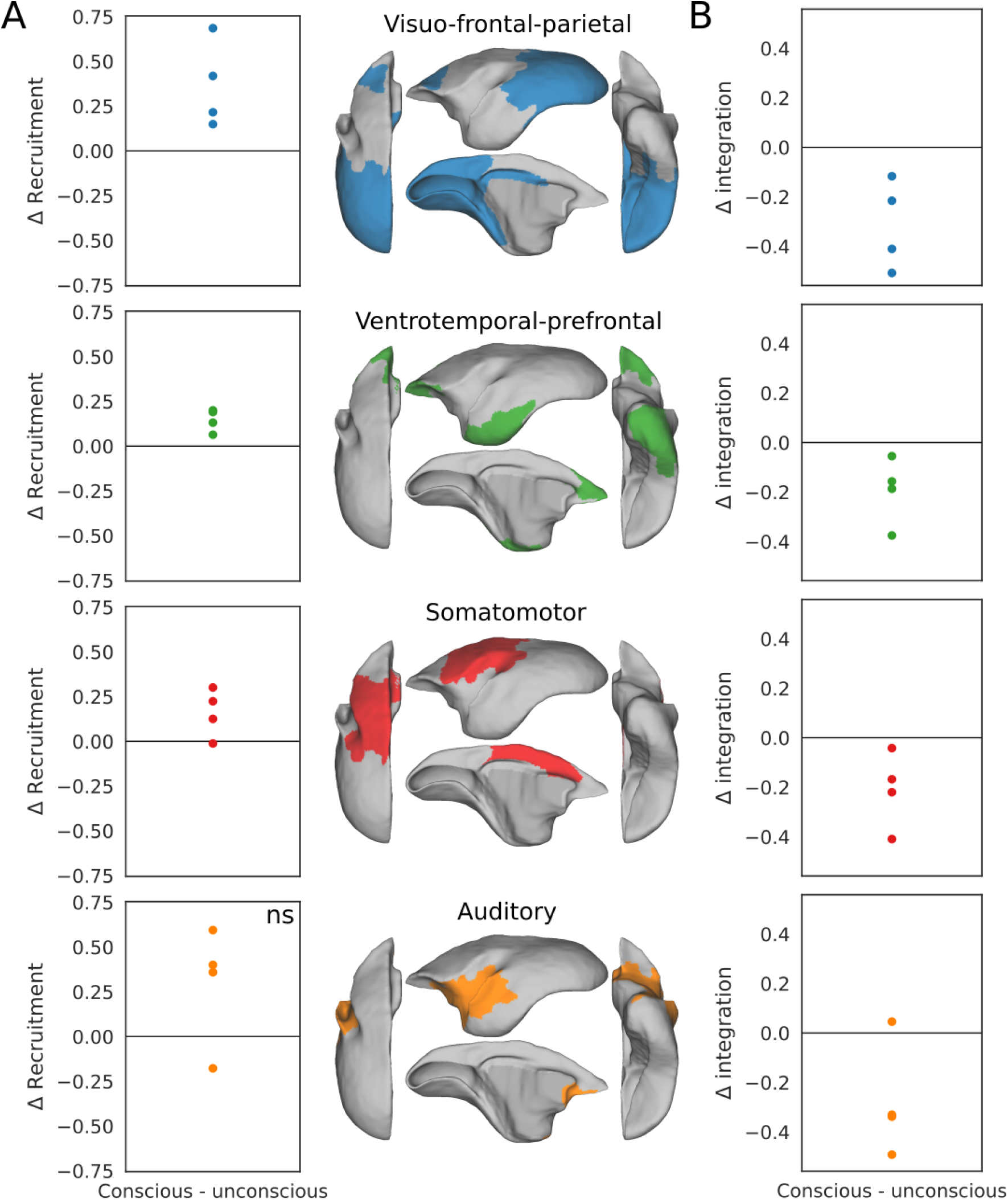
Subnetworks derived from dynamic modules were more distinct during consciousness than during unconsciousness **(A)** Recruitment of the visuo-fronto-parietal subnetwork [blue, t(3) = 3.519, p = 0.011], ventrotemporal-prefrontal [green, t(3) = 5.373, p = 0.007] and somatomotor [red, t(3) = 2.730, p = 0.038] subnetworks was significantly stronger in the awake condition than the anaesthetic condition, but the condition-dependent difference in recruitment of the auditory subnetwork (orange) was non-significant [t(3) = 2.054, p = 0.081], indicated by ‘ns’ in the bottom panel. **B** Each subnetwork was more integrated with the other three subnetworks in the anaesthetic condition [visuo-front-parietal: t(3) = −4.052, p = 0.009; ventrotemporal-prefrontal: t(3) = −3.341 p = 0.024; somatomotor: t(3) = −3.173, p = 0.018; auditory: t(3) = −2.813, p = 0.035]. As above, ‘Δ’ refers to the difference between conditions (awake minus anaesthetic).

#### Subnetwork recruitment and integration

Using the four consensus subnetworks derived by clustering the module allegiance matrix, we quantified the interaction within and between these subnetworks over time following the approach by Bassett et al. 17. Accordingly, the interaction between any two subnetworks, *k*_1_ and *k*_2_, was defined as 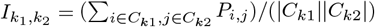, where *C*_*k*1_ and *C*_*k*2_ are the sets of regions belonging to subnetworks *k*_1_ and *k*_2_ respectively, |*C*_*k*_| is the number of regions in subnetwork *k*, and *P*_*i,j*_ is the module allegiance between regions *i* and *j. Recruitment* of a subnetwork *k* refers to the interaction of that subnetwork with itself (*I*_*k,k*_), calculated by setting *k*_1_ = *k*_2_ = *k*. Recruitment captures the stability or persistence of a subnetwork’s membership over time. *Integration* between two distinct subnetworks *k*_1_ and *k*_2_ (*k*_1_ ≠ *k*_2_) is defined as the normalized interaction between them: 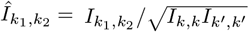. Integration quantifies the degree of interaction or ‘cross-talk’ between different subnetworks. For each subject, these metrics were calculated from its module allegiance matrix in each condition.

#### Statistical analysis

We used non-parametric permutation testing to assess statistical significance for all comparisons between the awake and anaesthetic conditions. This approach makes minimal assumptions about the distribution of the data. For each condition-dependent difference metric *X* (FC-SC distance, node strength, number of modules, flexibility, recruitment and integration), we calculated the paired t-statistic [*t*(3) = *µ*_X_*/*(*σ*_X_*/* 4)] across the four subjects based on the mean difference between conditions (awake minus anaesthetic). To generate the null distribution, we randomly permuted the condition labels (awake, anaesthetic) for the six scans within each subject 10,000 times. For each permutation, we recalculated the mean value of the metric for the randomly labelled ‘awake’ and ‘anaesthetic’ conditions for that subject, computed the difference, and then calculated the tstatistic across the four subjects based on these permuted differences. This procedure yielded a null distribution of 10,000 t-statistics under the hypothesis that there was no difference between conditions. The actual t-statistic calculated from the unpermuted data was then compared to this null distribution. A difference between conditions was considered statistically significant if the actual t-statistic fell in the extreme 5% of the null distribution. This procedure was applied independently for all cases of permutation testing.

## ACKNOWLEDGEMENTS

This work was supported by the Canadian Institutes of Health Research and the Natural Sciences and Engineering Research Council of Canada.

